# Experience-dependent learning and myelin plasticity in individuals with stroke

**DOI:** 10.1101/2022.02.17.480894

**Authors:** Cristina Rubino, Beverley C. Larssen, Larissa K. Chiu, Hanwen Liu, Sarah N. Kraeutner, Niruthikha Mahendran, Ronan Denyer, Bimal Lakhani, Michael R. Borich, Cornelia Laule, Lara A. Boyd

## Abstract

**Background:** Injury to brain myelin disrupts motor performance and learning, however it is not clear if myelination is modulated by skilled motor practice or by recovery after stroke. Multi-component relaxation imaging can be used to measure water trapped between myelin bilayers which is expressed as myelin water fraction.

The purpose of this study was to examine the effect of experience-dependent learning on myelin plasticity using multi-component relaxation imaging in individuals with stroke.

**Methods:** Thirty-two individuals with chronic stroke (>6 months) and twenty-seven healthy controls completed 4 weeks of skilled motor practice using a complex, gamified reaching task. Multi-component relaxation imaging-derived myelin water fraction was obtained before and after training. Seven brain regions associated with motor learning and sensorimotor function were investigated.

**Results:** All participants improved task-specific reaching movements after training. In individuals with stroke: 1) pre-training myelin water fraction was lower in motor brain regions but higher in the cingulum compared to controls, 2) pre-training myelin water fraction in motor and sensorimotor regions was positively associated with learning rate, and 3) myelin water fraction was increased in the ipsilesional (contralateral to the trained arm) superior longitudinal fasciculus following skilled motor practice.

**Conclusions:** Findings indicate that after stroke, myelin water fraction is related to measures of motor learning and modulated by 4 weeks of skilled motor practice with the paretic limb. Myelin water fraction can be enhanced in the chronic stage of stroke and may be an important target for upper-limb motor recovery.

## Introduction

Damage to the brain from stroke can lead to sensory, motor, and cognitive dysfunction.^1–3^ One of the most common and debilitating consequences of stroke is hemiparesis.^4–6^ There can be limits to motor recovery achieved as a result of both spontaneous biological recovery processes and in response to therapeutic intervention. Indeed, more than 40% of individuals are left with moderate to severe limb motor impairment,^7^ which impacts independence and ability to perform daily activities.^8^ Thus, increased understanding of motor learning, and associated changes in brain structure and function is crucial to enhancing rehabilitative strategies and recovery from stroke.

Changes in brain white matter may be an important component of motor recovery following stroke.^9–13^ Studies of stroke recovery biomarkers demonstrate that neuroimaging measures of white matter can predict long-term functional outcomes. Specifically, impairement in the corticospinal tract (CST), as measured by diffusion tensor imaging (DTI), is associated with poorer clinical outcomes (i.e., assessed using the FM-UE, Motricity Index, National Institute of health Stroke Scale, Functional Independence Measure) in the acute as well as chronic stage of stroke.^6,11,14–18^ This information may be used in combination with rehabilitation interventions to promote stroke recovery. Furthermore, while DTI data are useful in grossly characterizing white matter, multiple structural features may contribute to the quantitative metrics describing diffusion behaviour, including: 1) axonal membrane status, 2) myelin sheath thickness, 3) number of intracellular neurofilaments and microtubules, 4) axonal packing density, 5) inflammation and edema and 6) crossing fibres.^19,20^ Traditional DTI techniques assign one unique fiber orientation within each brain voxel. However, multiple fiber tracts in different orientations are present in more than 90% of brain white matter.^21^ Therefore, it remains unclear how DTI-derived metrics, such as fractional anisotropy (FA), specifically relate to white matter tissue composition.^20,22^ White matter imaging instead is a more specific measure of white matter and may be an important technique in advancing our understanding of how stroke disrupts brain myelin.

Myelinated axons are a major component of white matter and are crucial for fast and efficient information transfer in the brain.^23–25^ While impairment of myelination is associated with learning deficits, motor and cognitive dysfunction, and aging,^26–30^ myelin formation and regulation may be modified by experience-dependent motor learning.^29–36^ Animal model studies showed that blocking the formation of new oligodendrocytes (cells responsible for producing myelin) hindered complex wheel running,^29,30^ and skilled training enhances brain myelin in healthy mice.^32^ Additionally, motor learning following injury promoted remyelination and enhanced oligodendrogenesis.^33,34^

While it is clear that impairment of myelination disrupts motor performance and learning,^29,30,32–34^ examples of change in myelin associated with skilled motor practice in humans have been scarce. In part, this stems from technological limitations that until recently, limited the mapping of myelin in humans *in vivo*. Advances in MRI now allow for whole brain myelin mapping via multi-component relaxation imaging (MCRI). MCRI in the human brain can separate the magnetic resonance proton signal T_2_ relaxation time into three components: 1) a long T_2_ component (> 2 s) that is attributed to cerebral spinal fluid, 2) an intermediate T_2_ component (∼80 ms) attributed to intra/extracellular water, and 3) a short T_2_ component (< 40 ms) that reflects water trapped between myelin bilayers (myelin water).^37,38^ The amount of myelin water relative to the total water is termed the myelin water fraction (MWF) and validation studies in animal and human models confirm that MWF is an *in vivo* marker for myelin content in the brain.^39^ Formalin-fixed human brains yield T_2_ distributions similar to those found *in vivo*, and histopathological studies show strong correlations between MWF and staining for myelin.^39–41^

Myelin water imaging has been used to study the brain and spinal cord in many myelin-related diseases.^42^ For example, studies of individuals with multiple sclerosis reveal decreased MWF in both lesions and normally appearing white matter,^43,44^ and multicomponent driven equilibrium single pulse observation of T_1_ and T_2_ determined MWF is related to severity of clinical signs.^45^ After stroke MWF is decreased in the bilateral posterior limbs of the internal capsule, and across the whole cerebrum, compared to healthy controls.^46^ These data illustrate that stroke impairs myelin even in sites that are distant from the lesion. Even though animal model studies show that motor learning can promote remyelination, it is not clear whether skilled motor training can modify myelin in the stroke-damaged brain. To date, no work has examined training-induced change in myelin in individual brain regions after stroke.

The current study used MCRI to map myelin before and after skilled motor practice in individuals with chronic stroke in three motor and four sensorimotor regions of interest (ROIs) of the ipsilesional, trained hemisphere. Primary motor regions (including, premotor area, primary motor cortex and basal ganglia) are important for limb motor function. After stroke, functional connectivity shifts to nearby networks not immediately impacted by the lesion.^47^ Though the corticospinal tract may be important for motor recovery,^18,48^ it is likely that individuals with damage to the CST who are in the chronic stage of stroke recovery (>6 months post) employ alternate pathways to support limb motor function. Therefore, in the current study of individuals with chronic stroke, we broadened the inclusion criteria to cover regions that could possibly be affected by the stroke beyond those traditionally studied, including association fibers and sensorimotor regions.

The objectives of our study were to: 1) examine the effect of stroke on human brain myelin compared to healthy individuals, 2) determine the relationship between baseline myelin and motor skill learning of a gamified sensorimotor reaching task, and 3) assess changes in myelin following skilled motor practice in individuals with stroke and controls. We hypothesized that we would find: 1) decreased brain myelin, as indexed by MWF, in the lesioned hemisphere of individuals with stroke compared to the non-dominant hemisphere in controls, 2) a positive relationship between pre-training MWF and rate of motor skill acquisition in individuals with stroke as well as controls, and 3) increased MWF immediately following skilled motor practice in the ipsilesional, trained hemisphere of individuals with stroke and non-dominant hemisphere of controls.

## Materials and methods

### Participants

Forty-two individuals with chronic (>6 months) stroke and thirty-one healthy individuals were recruited for this study. Individuals in the stroke group were included if they had a middle cerebral artery infarct at least six months prior to enrollment and were between the ages of 35-85 years old. Individuals from both groups were excluded if they: 1) were unable to complete MRI scanning, 2) showed signs of dementia (a score of <23 on the Montreal Cognitive Assessment^49^), or 3) had history of head trauma, seizure, psychiatric diagnosis, neurodegenerative disorder, substance abuse, or other neurological or muscular deficits that affected vision or manual control. All testing was conducted at the University of British Columbia. Consent from each participant was obtained according to the Declaration of Helsinki. The Research Ethics Board at the University of British Columbia approved all study procedures.

### Clinical Assessment

A trained physical therapist or clinical student in-training rated motor impairment of individuals with stroke using the motor portion of the Fugl-Meyer upper extremity (FM-UE) scale^50^ 24 hours before motor skill practice. The FM-UE is a measure of upper extremity impairment and is rated on a scale from 0-66, with higher scores reflecting less impairment.^50^ The FM-UE assessment was used in the current study to characterize the spread of stroke severity in the sample.

### Behavioural task

#### Apparatus

All participants engaged in a semi-immersive virtual reality-based intercept and release task (<u>TR</u>ack <u>A</u>nd <u>I</u>ntercept <u>T</u>ask; TRAIT; details reported in.^35^ Briefly, TRAIT was presented on a 46-inch monitor, viewed at 72 inches away (screen refresh rate 59 Hz). A Microsoft Kinect (model no. 1517, Kinect for Windows; Microsoft, Redmond, WA) camera was used for motion detection and communication between the participant and the computer task. Before each session, the Microsoft Kinect was calibrated using a four-point grid; this enabled the workspace to be personalized for each individual in every session and allowed participation from individuals with varying degrees of range of motion. The interactive environment of the task was set in “outer-space”, where the position of the hand, object, and target were coded as a spaceship, asteroid, and sun, respectively. The cartesian (X,Y,Z) coordinates of the hand’s center-point were captured by the Microsoft Kinect sensor. The task was created to induce a large dose of skilled movements while maintaining motivation and engagement throughout practice.^52^

#### Procedure

A total of 10 sessions of skilled motor practice were completed over 4 weeks (Fig. 1). Participants were asked to “save the world” by controlling an on-screen icon (spaceship) using movements of the relevant arm (stroke-affected arm for stroke participants or non-dominant arm for control participants) to intercept a virtual moving object (asteroid) as it emerged from the side of the screen. Once intercepted, participants had to accurately throw the asteroid so that it hit a target (the sun), causing the asteroid to explode. The location of the target varied randomly on the screen. Auditory (i.e., sound effects) and visual feedback, (i.e., asteroid exploding) as well as knowledge of results (i.e., a numerical score provided at the end of each trial and block) were used to maintain motivation and engagement. Participants were instructed to complete each trial of the task as “quickly and accurately as possible”.

**Figure 1.**
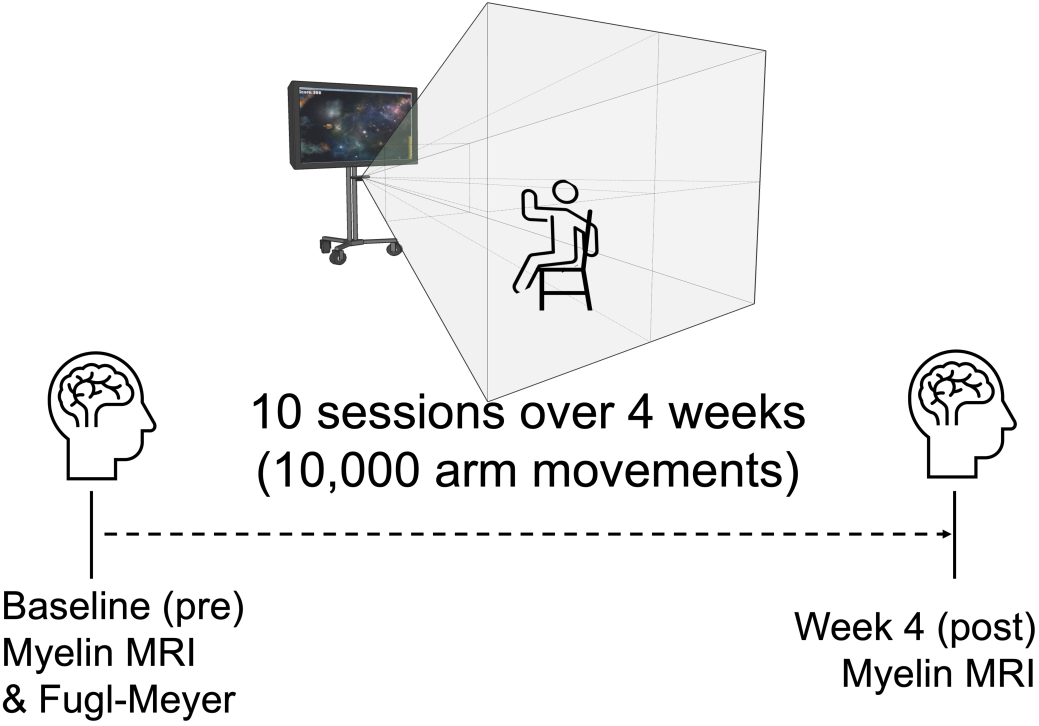
Timeline of the experimental design. Participants engaged in ten sessions of a gamified sensorimotor reaching task over four weeks, amounting to 10,000 total arm movements. Participants used their affected (stroke) or non-dominant arm (control). Within 24 hours prior to and following training, participants underwent MRI to capture changes in myelin water fraction (MWF).

During each session participants performed 5 blocks of the task, lasting approximately 45 mins. Each block contained 200 movements (100 object interceptions and 100 object throws), for a total of 1000 skilled arm movements per session and 10,000 total skilled movements across the experiment. Task difficulty was manipulated by increasing the speed of the object, decreasing the size of the object or decreasing the size of the target. Participants progressed through levels of increasing task difficulty as their skill improved. The skill criterion to advance to the next level of task difficulty was an 80% success rate of object interception and hitting the target on two consecutive practice blocks. The task was designed to challenge a wide range of participants, maintain engagement, and prevent plateaus in performance. The number of practice days and trials were determined from our prior work^35^ based on the principle that both repetition and intensity are important to stimulate experience-dependent neuroplastic change.^53^

### MRI Acquisition

MRI data were acquired on a 3.0T Philips Achieva (Best, The Netherlands) whole body MRI scanner using an eight-channel sensitivity encoding head coil and parallel imaging. MR experiments included: (1) 3D T_1_ turbo field echo for anatomical identification (TE/TR = 3.6/7.4 ms, flip angle θ = 6°, field of view (FOV) = 256 × 256 mm^2^, 160 slices, 1 mm slice thickness, scan time = 3.2 min); and (2) whole-cerebrum 32-echo three-dimensional Gradient and Spin Echo (GRASE) for T_2_ measurement (TE/TR = 10/1000 ms, 20 slices acquired at 5 mm slice thickness, 40 slices reconstructed at 2.5 mm slice thickness (*i.e*. zero filled interpolation), slice oversampling factor = 1.3 (*i.e*. 26 slices were actually acquired but only the central 20 were reconstructed), in-plane voxel size = 1 × 1 mm^2^, SENSE = 2, 232 × 192 matrix, receiver bandwidth = 188 kHz, axial orientation, acquisition time = 14.4 min).^51^ T_1_-weighted and GRASE sequences were obtained in the same session. Participants were scanned at two time points, within 24 hours before motor training began and after completing four weeks of motor practice.

### Data analyses

We quantified motor skill acquisition by exponentially fitting object interception time for each successful trial over training, using equation 1. Using information from all successful trials, we calculated, the rate of skill acquisition, change in movement time, and movement time at plateau in performance.^54^

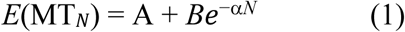

*E*(MT) is the expected value of movement time (MT) on trial *N*. **A** (seconds) defines the movement time at which the participant plateaued in performance. **α** (seconds/trial; our primary outcome measure) quantifies the rate of skill acquisition to the point of plateau, and ***B*** (seconds) is a measure of overall change in movement time from the beginning of training to the point of performance plateau.

### MWF Map Generation

Myelin water fraction (MWF) maps (Fig. 2A) were generated for each individual. Using the 32-echo GRASE data, voxel-wise T_2_ distributions were calculated using a modified Extended Phase Graph algorithm combined with regularized non-negative least squares and flip angle optimization (in-house MATLAB software - requested here: https://mriresearch.med.ubc.ca/news-projects/myelin-water-fraction/).^51,55^ MWF was defined as the sum of the amplitudes within a short T_2_ signal (15-40 ms) divided by the sum of the amplitudes for the total T_2_ distribution. Representative MWF maps from one control and stroke participant are shown in Fig. 2A.

**Figure 2.**
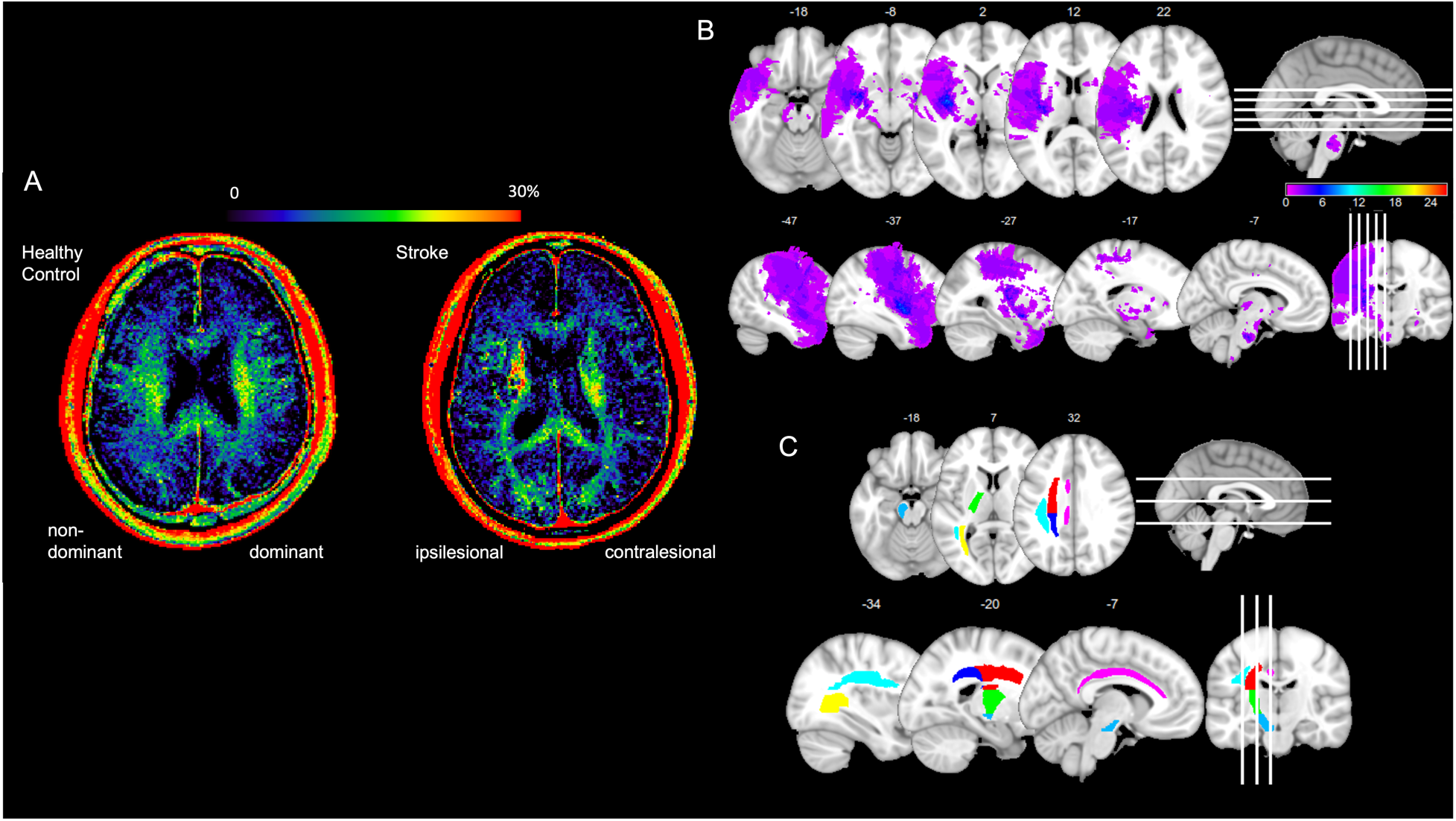
Myelin water fraction map, stroke lesion map and atlas regions. (**A**) Representative slice of the mean MWF atlas for one control and one stroke participant. Colour scale indicates percentage of myelin water fraction in each voxel. (**B**) Overlay map of individual stroke lesions of all 25 participants included in the final analyses. Maps are overlaid on a MNI 1mm brain. All lesions were flipped to the right hemisphere. Colour scale indicates the number of stroke participants having a lesion that region. Stroke lesions are distributed across regions supported by the middle cerebral artery. (**C**) JHU-ICBM-DTI-81 atlas regions of the cerebral peduncles (light blue), posterior limb of internal capsule (green), superior corona radiata (red), posterior corona radiata (blue), posterior thalamic radiations (yellow), superior longitudinal fasciculus (cyan) and cingulum-cingulate gyrus (violet).

### Lesion Masking

Lesion masking was performed manually using T_1_ scans by two trained individuals. Masks were used to exclude lesioned tissue from analyses and to create a lesion heat map (Fig. 2B). Masks were created in individual T_1_ space using the tool mode in FSLeyes (https://fsl.fmrib.ox.ac.uk/fsl/fslwiki/FSLeyes). All lesioned regions in the brain of at least 2 voxels in size were masked. Masks drawn at the pre time point were warped to T_1_ space at the post time point and visually inspected for consistency.

### Regions of Interest

The John Hopkins University International Consortium of Brain Mapping (JHU-ICBM) DTI-81 white-matter labels atlas^56^ was used to identify subcortical regions of interest (ROIs). ROIs were selected a priori based on their function in motor control and learning as well as sensorimotor integration,^57–61^ and included <u>3 motor regions</u> (cerebral peduncles (CP), posterior limb of the internal capsule (PLIC), superior corona radiata (SCR)) and <u>4 sensorimotor regions</u> (posterior thalamic radiations (PTR), posterior corona radiata (PCR), cingulate gyrus-cingulum (CCG), superior longitudinal fasciculus (SLF))(Fig. 2C).

### MWF Extraction

To extract MWF from ROIs for all individuals, several processing steps were conducted using FSL command line tools (FMRIB Software Library, Oxford).^62^ This included, transforming atlas ROIs from MNI space, and lesion masks from T_1_ space (stroke group only), to individual native space. The specific processing steps are outlined below:

1. GRASE to T_1_: First echo of the GRASE scans were linearly registered (12 degrees of freedom) to their respective T_1_ scans.
2. T_1_ to MNI: For stroke scans, individual lesion masks were overlaid onto their respective T_1_ images to improve brain extraction (skull-stripping) and registration outcome. This was done by assigning a value to the lesion mask that was similar to the average tissue intensity value in each individual’s brain. Lesion masks were removed following successful brain extraction and registration (and later used in Step 5). For all scans, brain extracted T_1_ scans were then non-linearly warped to 1mm MNI space.
3. MNI to GRASE: Warping fields from steps 1 & 2 were inverted to create a warping field of MNI to GRASE space. The warp was used to transform ROIs in 1 mm MNI space to native GRASE space.
4. Lesion masks in T_1_ to GRASE: Inverted linear registrations from step 1 transformed lesion masks (in T_1_ space) to GRASE space.
5. Extraction of mean MWF: For stroke scans, lesion masks were subtracted from ROIs that overlapped with the stroke lesion. For all scans, mean MWF for each ROI in native space was extracted and used in statistical analyses.

Scans were visually inspected following each processing step to ensure manipulations were successful. In addition, we quantified alignment of ROIs into native space by calculating a coefficient of variation (COV: mean MWF divided by its standard deviation) for each ROI. High COV values (i.e., larger than 0.75) are likely produced by misalignment and poor registration.^63^ All ROIs had a COV of less than 0.75.

### Statistical analysis

All dependent measures were assessed for normality and homogeneity prior to performing parametric statistical testing. Non-parametric testing was conducted for variables that did not meet assumptions of normality and homogeneity. Furthermore, paired-sample *t*-tests were performed on ROI volumes (pre versus post) to ensure consistency across time points and control for influence of ROI volume on mean MWF. All statistical analyses were conducted using SPSS v26 (IBM SPSS Statistics) with *P* < 0.05 set *a priori* to denote statistical significance.

To address our research questions, we performed the following statistical tests:

1. Independent samples *t*-tests were conducted using ipsilesional (stroke)/non-dominant (controls) mean MWF of each ROI as the dependent variable to test the difference between individuals with stroke and controls before training. Bonferroni correction for multiple ROI comparisons was made.
2. Bivariate correlations were conducted to test the relationship between ipsilesional (stroke)/non-dominant (controls) mean MWF before training and rate of change of motor skill acquisition (as measured by the α derived from Equation 1).
3. Multiple one-way repeated measures ANOVA were conducted for each ROI to test change in mean MWF before and after training in the trained hemisphere (ipsilesional hemisphere in stroke participants or non-dominant hemisphere in control participants). Bonferroni correction for multiple ROI comparisons was made.

## Results

25 individuals with stroke (66.6±10.6 years old, 8 female, 59.4±45.5 months since stroke, 15 right-affected hemisphere, 18-66 FM-UE score range) and 24 controls (64.5±8.4 years old, 17 female, 22 right-handed) were included in the final analyses. Ten stroke and four control participants dropped out after the first MRI. Data from seven stroke and three control participants were excluded for image registration issues involving registration of T1 to MNI or MNI to GRASE space, or behavioural data >3SD away from the mean. Age was not significantly different between groups (t(47)= -0.737, *P* = 0.465). Sex was significantly different between groups (t(45) = 2.89, *P* = 0.005) where the stroke group had higher proportion of male participants compared to the control group. Demographic information (age, sex, time since stroke, FM-UE score, affected hemisphere and lesion location) for each individual included in the final analyses are reported in Table 1.

**Table 1.** Demographic information of stroke and control participants included in the final analyses.

| Stroke |  |  |  |  |  |  | Control |  |  |
| --- | --- | --- | --- | --- | --- | --- | --- | --- | --- |
| Participant | Age | Sex | Time since stroke (months) | Fugl-Meyer (UE) score | Affected hemisphere | Lesion location | Participant | Age | Sex |
| S01 | 57 | F | 162 | 32 | R | Middle frontal gyrus | C01 | 67 | M |
| S02 | 51 | F | 47 | 29 | R | Basal ganglia | C02 | 63 | F |
| S03 | 58 | M | 96 | 59 | L | Inferior frontal gyrus | C03 | 81 | M |
| S04 | 72 | M | 49 | 50 | R | Basal ganglia | C04 | 58 | F |
| S05 | 71 | M | 117 | 59 | R | Middle frontal gyrus | C05 | 74 | F |
| S06 | 77 | M | 32 | 58 | L | Pons | C06 | 62 | F |
| S07 | 70 | F | 76 | 64 | R | Corticospinal tract | C07 | 62 | F |
| S08 | 60 | M | 188 | 31 | R | Parietal lobe | C08 | 61 | F |
| S09 | 73 | M | 60 | 54 | L | Thalamus | C09 | 63 | M |
| S10 | 69 | M | 66 | 51 | L | Pons | C10 | 73 | F |
| S11 | 66 | M | 10 | 66 | R | Putamen | C11 | 67 | F |
| S12 | 37 | F | 84 | 18 | L | Basal ganglia | C12 | 75 | M |
| S13 | 79 | M | 27 | 54 | R | Basal ganglia | C13 | 71 | M |
| S14 | 62 | M | 8 | 59 | R | Putamen | C14 | 65 | F |
| S15 | 80 | M | 40 | 62 | R | Basal ganglia | C15 | 77 | F |
| S16 | 78 | F | 28 | 64 | R | Pons | C16 | 63 | F |
| S17 | 75 | M | 41 | 65 | L | Basal ganglia | C17 | 58 | F |
| S18 | 79 | F | 51 | 59 | R | Basal ganglia | C18 | 68 | M |
| S19 | 59 | M | 61 | 33 | L | Post central gyrus | C19 | 58 | M |
| S20 | 61 | M | 62 | 28 | L | Insular cortex | C20 | 56 | F |
| S21 | 71 | F | 94 | 52 | L | Basal ganglia | C21 | 47 | F |
| S22 | 74 | F | 19 | 39 | R | Precentral gyrus | C22 | 50 | F |
| S23 | 51 | M | 14 | 23 | R | Corticospinal tract | C23 | 58 | F |
| S24 | 73 | M | 47 | 64 | R | Thalamus | C24 | 73 | F |
| S25 | 62 | M | 6 | 62 | L | Amygdala |  |  |  |

### Motor skill practice

An average of 84.2±7.9% (stroke) and 85.9±5.6% (controls) of the asteroids were successfully intercepted across all levels of task difficulty where individuals in the stroke group reached level 7 (range: 1 – 11) and the control group reached level 9 (range: 6 – 12) of 12 total levels. All participants learned the motor task (faster movement time at the end compared to the beginning of training; Supplementary Fig. 1A, B). Exponential curve fitting using Equation 1 [*R*^2^=0.29, range=0.16-0.87±0.48 (stroke); *R*^2^=0.55, range=0.24-0.83±0.24 (control)] revealed the following findings. The average trial number at which individuals plateaued in movement time was 3169.9±618.8 (stroke) and 3080.3±809 (control) of 5,000 total trials. Average trial numbers were not significantly different between groups (t(47) = 0.615, *P* = 0.54). Resultant mean ***B* values**, magnitude change in movement time from training plateau to beginning of training, were 1.8±0.59 seconds (stroke) and 1.41±0.38 seconds (controls). Mean *B* values were significantly different between groups (t_*B*_(47) = 2.66, *P* = 0.01). **α values**, rate of reduction of movement time per trial (learning rate; our primary behavioural outcome measure), were 0.0009±0.00018 seconds/trial (stroke) and 0.001±0.00025 seconds/trial (controls). Mean α values were not significantly different between groups (t_α_(47) = -1.43, *P* = 0.16). **A values**, movement time at training plateau, were: 1.09±0.47 seconds (stroke) and 0.76±0.06 seconds (controls). Mean A values were significantly different between groups (t_A_(47) = 3.84, *P* = 0.0003). Shapiro-Wilk and Levene’s tests showed that all behavioural measures met assumptions of normality and homogeneity.

### Myelin water fraction

Shapiro-Wilk and Levene’s tests showed mean MWF of each ROI per group met assumptions of normality and homogeneity except the CP, PLIC, and SCR. Analyses of the three ROIs that did not meet assumptions of homogeneity were conducted with a Kruskal-Wallis rank sum test. Age (years) and time since stroke (months) were not significantly correlated with mean MWF of any ROI. Volume of ROIs before and after training were not significantly different within individuals.

### Aim 1: Baseline MWF between groups

Independent samples *t*-tests as well as Kruskal-Wallis rank sum tests for CP, PLIC and SCR revealed significant group differences in baseline (pre-training) mean MWF (Fig. 3) for all ROIs, except CP and PCR. Lower MWF in SCR, PLIC, PTR and SLF was observed in the stroke ipsilesional hemisphere compared to controls’ non-dominant hemisphere, while higher MWF in the CCG was observed in the stroke ipsilesional hemisphere compared to controls’ non-dominant hemisphere (*P* < 0.05, uncorrected). After Bonferroni correction, SCR and PLIC MWF remained significantly lower in the stroke ipsilesional hemisphere compared to controls’ non-dominant hemisphere (*P* < 0.007). Mean differences and *P*-values for each ROI are summarized in Table 2.

**Table 2.** Mean MWF differences at baseline for each ROI.

| ROI | Stroke<br>mean MWF<br>(SD) | Control<br>mean MWF<br>(SD) | % difference<sup>a</sup> | P-value |
| --- | --- | --- | --- | --- |
| Cerebral peduncles | 0.240 (0.066) | 0.246 (0.035) | 0.5 | 0.410 |
| Superior corona radiata | 0.103 (0.027) | 0.135 (0.017) | 3.2 | **<0.001<sup>b\*</sup>** |
| Posterior limb internal capsule | 0.158 (0.045) | 0.186 (0.026) | 2.7 | **0.005<sup>b\*</sup>** |
| Posterior thalamic radiations | 0.110 (0.023) | 0.126 (0.016) | 1.6 | 0.007* |
| Posterior corona radiata | 0.099 (0.021) | 0.106 (0.015) | 0.7 | 0.211 |
| Cingulum-cingulate gyrus | 0.081 (0.023) | 0.068 (0.016) | 1.3 | 0.028* |
| Superior longitudinal fasciculus | 0.112 (0.024) | 0.126 (0.016) | 1.4 | 0.023* |

\* $p < 0.05$
<sup>a</sup>Percent (%) difference = $[(\text{meanMWF}_{\text{stroke}} - \text{meanMWF}_{\text{control}}) * 100]$
<sup>b</sup>Bolded values indicate statistical significance was reached after Bonferroni correction ( $P < 0.007$ )

**Figure 3.**
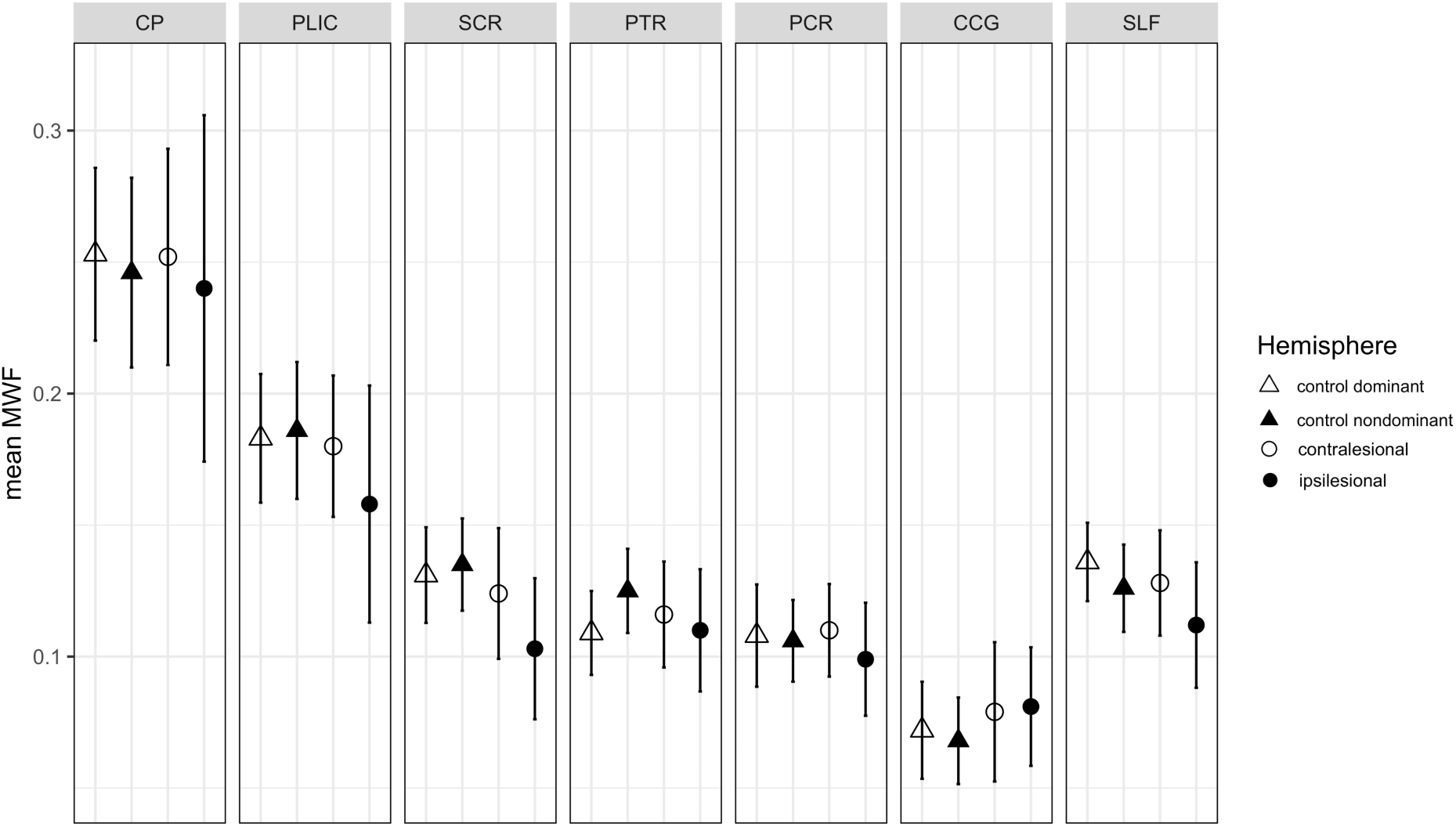
Myelin water fraction between stroke and control participants. Mean and standard deviation of MWF at baseline (prior to training) is plotted. Data are grouped by hemisphere, ipsilesional/contralesional (stroke) and dominant/non-dominant (control), and region of interest (ROI), CP: cerebral peduncles, PLIC: posterior limb of internal capsule, SCR: superior corona radiata, PTR: posterior thalamic radiations, PCR: posterior corona radiata, CCG: cingulum-cingulate gyrus, SLF: superior longitudinal fasciculus.

### Aim 2: Baseline MWF and motor skill acquisition

Pearson’s correlations revealed a significant positive relationship between pre-training MWF and rate of skill acquisition (α values; our primary outcome measure). In the stroke group, pre-training MWF in the ipsilesional CP (*r* = 0.5, *P* = 0.012), PLIC (*r* = 0.43, *P* = 0.033), SCR (*r* = 0.48, *P* = 0.016) and PCR (*r* = 0.5, *P* = 0.011; uncorrected) were significantly positively correlated with α values (Fig. 4). In the control group, pre-training MWF in the non-dominant PLIC (*r* = 0.49, *P* = 0.015; uncorrected) was significantly positively correlated with α values (Fig. 4). Correlation coefficients and *P*-values for each ROI are summarized in Supplementary Table 1. Exploratory correlation analyses of pre-training MWF and our secondary outcome measures (A and *B* values) revealed the following. There was a significant negative correlation between ipsilesional SCR (*r* = -0.41) and A values, but no significant result with *B* values in the stroke group. There was a significant positive relationship between non-dominant PLIC (*r* = 0.45, *P* < 0.05 uncorrected), SCR (*r* = 0.42), PCR (*r* = 0.45; all *P* < 0.05 uncorrected) and *B* values, but no significant result with A values in the control group.

**Figure 4.**
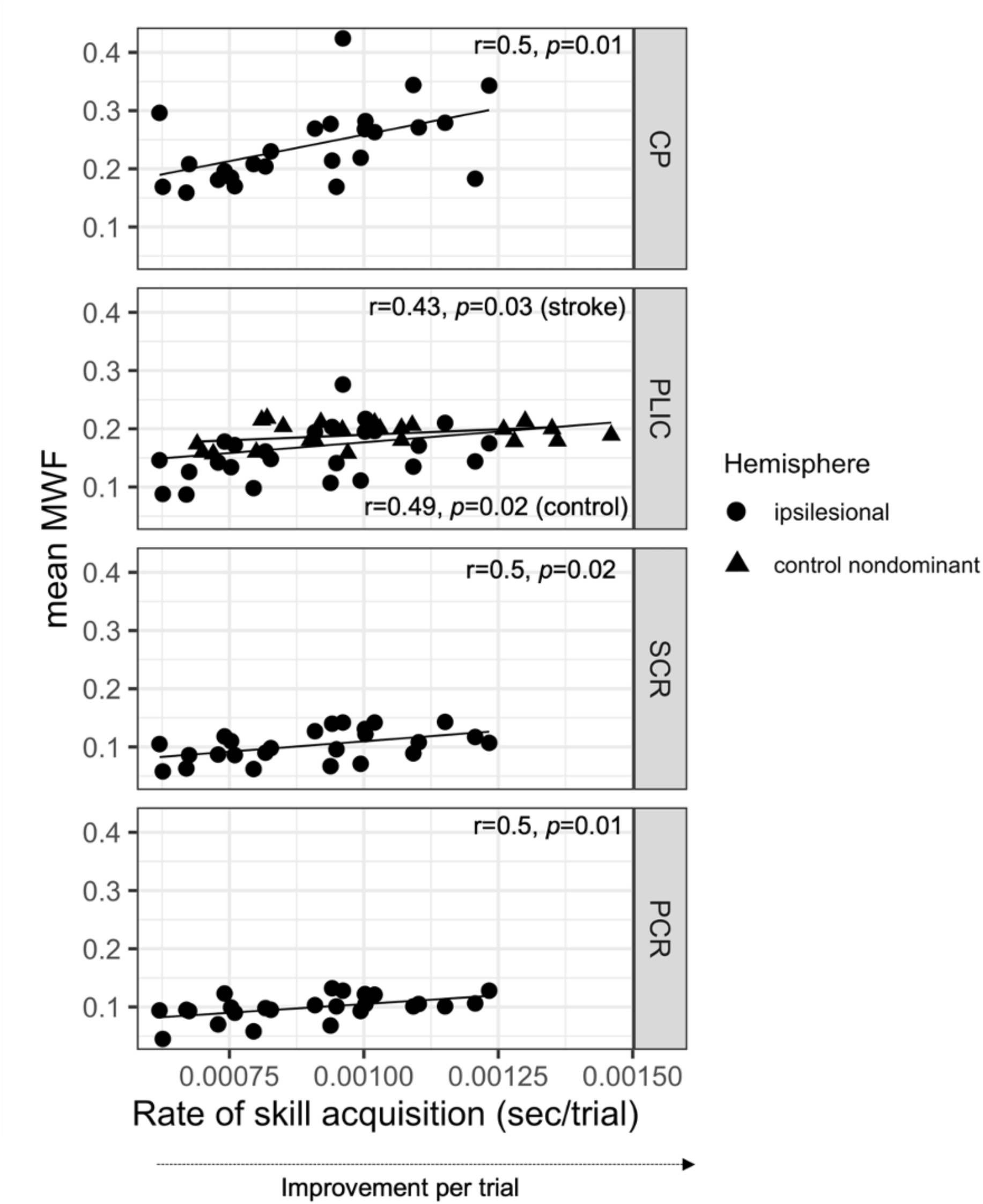
Correlation between mean MWF at baseline (prior to training) and rate of skill acquisition. Rate of skill acquisition is also referred to as α (derived from Equation 1) and represents improvement per trial. Statistically significant correlations (*P*<0.05) are grouped by hemisphere, ipsilesional (stroke) and non-dominant (control), and region of interest (ROI), CP: cerebral peduncles, PLIC: posterior limb of internal capsule, SCR: superior corona radiata, PCR: posterior corona radiata.

### Aim 3: Change in MWF after 5,000 trials of skilled motor practice

One-way repeated measures ANOVA’s (one for each ROI) in the stroke group revealed a significant effect of time for the SLF (Wilks’ Lambda = 0.8, F (1,24) = 6.1, *P* = 0.021; uncorrected) where mean MWF in ipsilesional SLF was greater post-compared to pre-training (Fig. 5). In separate one-way repeated measures ANOVA’s (one for each ROI) in the control group there was no significant effects observed between pre- and post-training mean MWF in the non-dominant hemisphere. A post-hoc *t*-test confirmed that mean MWF in the ipsilesional SLF at the post timepoint was no longer statistically different than mean MWF in the non-dominant SLF in controls. Mean MWF values per ROI, timepoint and group are summarized in Table 3.

**Table 3.**
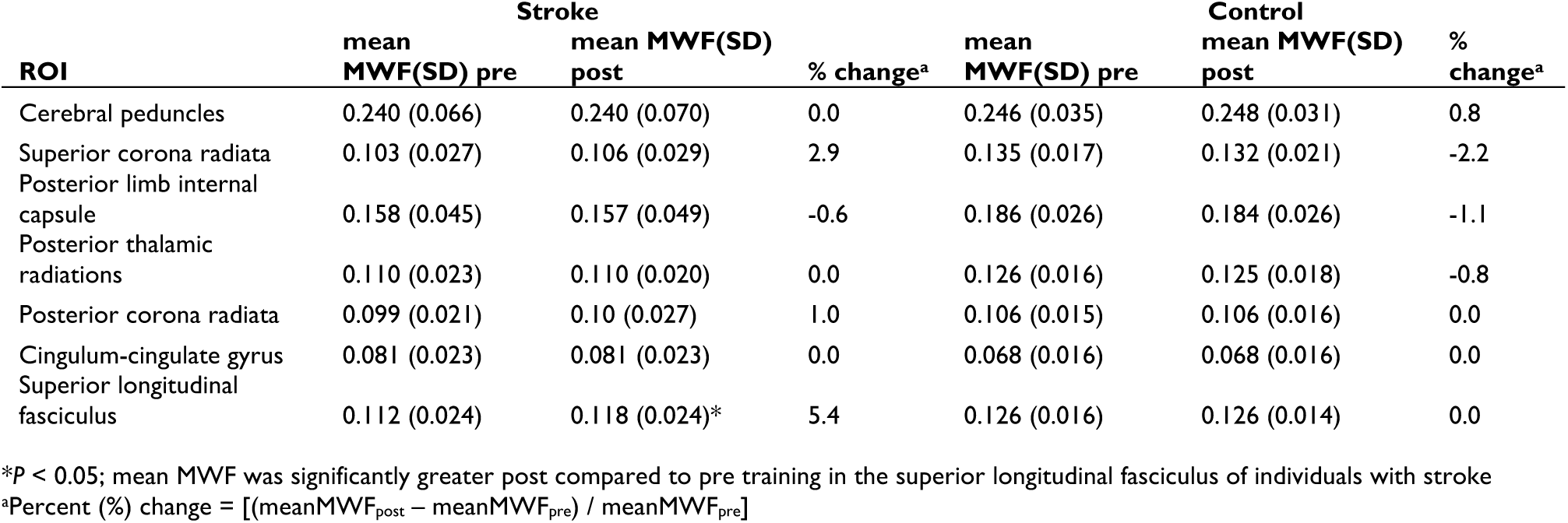
Mean MWF before and after training for each ROI.

**Figure 5.**
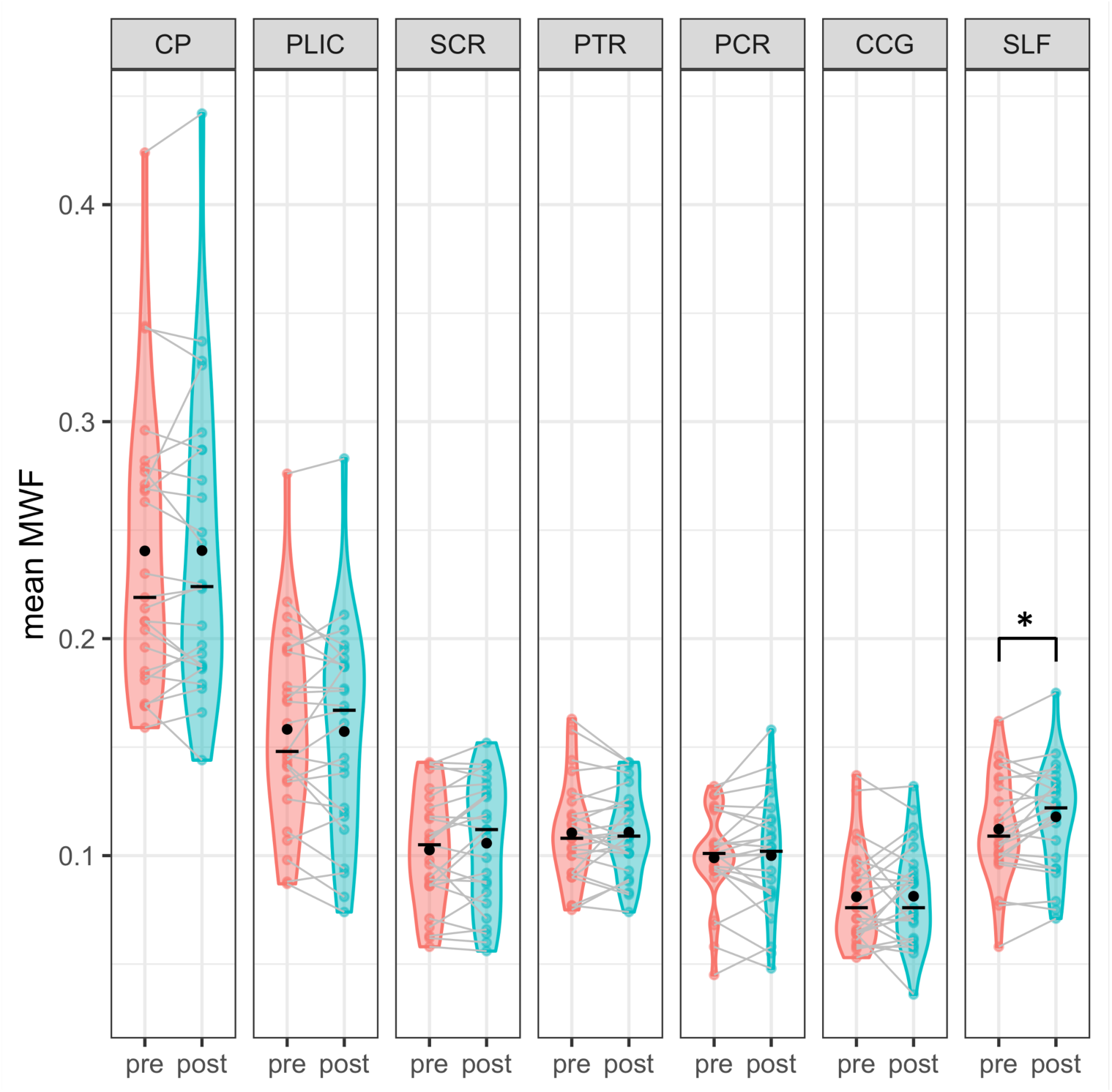
Change in mean MWF before and after training. Mean (black point), median (black line) and individual data points of the stroke ipsilesional (trained) hemisphere are plotted.

## Discussion

The current study examined how massed practice of skilled arm movement after stroke influenced myelin plasticity in the human brain. We compared MWF prior to and following 4 weeks of skilled motor practice in individuals with stroke and healthy controls. Consistent with our hypotheses, we observed: 1) decreased myelin water fraction in the ipsilesional hemisphere in individuals with stroke compared to healthy controls at baseline in all ROIs except the cingulum (which showed the opposite relationship), 2) a positive relationship between baseline myelin water fraction and learning rate, and 3) increased myelin water fraction following skilled motor practice in the hemisphere contralateral to the trained paretic arm in individuals with stroke. Contrary to our hypotheses, skilled motor practice did not significantly increase myelin water fraction in healthy controls. In summary, we provide the first evidence that skilled movement promotes myelin plasticity in the human stroke-injured brain.

### Modified MWF after stroke in motor and sensorimotor white matter regions

Our findings showed that myelination is changed in the stroke-injured hemisphere of individuals with chronic stroke compared to controls, within 5 white matter regions that contribute to visuomotor control. Aligned with prior work showing reduced MWF in motor projection pathways,^64^ pre-training MWF was reduced in ipsilesional PLIC and SCR compared to controls in the current study. While microstructural integrity of the pyramidal tract (primary descending motor fibers) is studied extensively after stroke and often associated with degree of motor impairment and recovery potential,^11,12,65–68^ other tracts may also be important markers for stroke recovery.^69,70^ Our data showed that myelination is altered within regions involved in sensorimotor function and remote to the infarct, including the PTR, SLF and cingulum. This is in line with wide-spread functional and structural network dysconnectivity findings from prior studies.^71–77^ Surprisingly, MWF was greater in the stroke ipsilesional cingulate gyrus compared to the homologous region in the contralesional hemisphere and in the controls (Fig. 3). Prior work showed that decreased activity in the cingulate is associated with aging,^78^ and increased functional recruitment was observed following high cognitive load in older adults.^79^

Furthermore, stroke results in increased activity between the default mode network and cingulate gyrus.^61^ Evidently, the cingulate gyrus serves an important role in memory, cognition and attention, and is disrupted in elderly healthy adults. Enhanced myelination in the cingulate gyrus after stroke may reflect a compensatory mechanism to support cognition, learning and memory following brain damage (refer to Bubb *et al*.^80^ for review of anatomical and functional organization of the cingulum). Critically, the current study provides evidence of wide-spread changes in myelination after stroke even in regions that were spared from direct lesion involvement.

### MWF is associated with learning rate of skilled motor practice

Here, we observed the relationship between myelination in four white matter regions and motor learning for individuals with stroke and controls. Our data showed that MWF in the PLIC is positively associated with learning rate of skilled motor practice in both individuals with stroke and healthy adults. Interestingly, additional regions involved in motor and sensorimotor function, such as the CP, SCR and PCR, were also positively associated with learning rate in the stroke but not in the control group (Fig. 4). Learning rate is the average rate of skill acquisition (improvement) per trial, such that a lower average rate (i.e., value closer to zero) indicates less improvement per trial compared to a higher average rate (i.e., value further from zero) which shows faster improvement. Our data suggest that higher MWF at baseline is associated with faster learning rate in stroke affected individuals. Furthermore, stroke participants whose motor behaviour changed at a faster rate and had more initial myelin also showed a faster initial performance. Together, movement times and MWF in the CP, PLIC, SCR, and PCR may help inform capacity for change during motor learning after stroke, however further analyses are needed to explicitly test each region’s unique contribution of MWF to motor learning.

Compared to controls, MWF was significantly reduced in the PLIC after stroke. This is consistent with other reports that show the importance of the PLIC for motor function^81^ and data showing that this tract is often damaged by stroke.^11,64,82^ Nonetheless motor learning occurs after stroke.^83,84^ Thus, it is likely that individuals compensate for stroke related damage by employing a distributed network of myelinated axons (beyond the PLIC). This idea is supported by prior work that showed that after stroke individuals rely on a dispersed functional network to learn as compared to age-matched controls.^47^ Both the current study of brain myelination, and our past work of functional connectivity,^47^ suggest that the motor cortex and regions in the parietal cortex interact to support motor learning after stroke. These interactions are not surprising given the need for parietal cortical function, which supports visuomotor integration during reaching movements, for motor task success.^85,86^

### Skilled motor practice enhances MWF after stroke

A primary objective of the current study was to examine experience-dependent neuroplastic change in white matter microstructure. Notably, research investigating white matter change in the brain after stroke, with a focus on myelination of axons (a critical structural component of white matter)^87^, is limited. This is the first study to examine changes in myelination (as measured by MWF) in the stroke damaged human brain, with advanced multi-component T_2_ relaxation imaging.^37^ Our results showed MWF increased in the SLF contralateral to the trained arm of individuals with stroke but not in controls following skilled motor practice (Fig. 5). Furthermore, a post-hoc analysis revealed that post-training MWF in the ipsilesional SLF was not different than the non-dominant hemisphere of controls, suggesting that myelination in the stroke damaged hemisphere may be enhanced after motor practice and comparable to MWF in controls. The SLF is involved in multiple functions, including working memory, language, visuospatial and motor function, and extends its fibers to frontal, parietal and temporal lobes (see Parlatini *et al*.^88^ for overview of SLF functional organization). Yet, the SLF is often overlooked, particularly as it relates to learning after stroke. Only two studies to date examined fronto-parietal involvement in chronic stroke motor performance. One showed that the integrity of the SLF is related to grasping imagery.^89^ Another revealed that individuals with extensive CST damage had greater functional connectivity in the ipsilesional fronto-parietal network associated with improved performance and higher fractional anisotropy of the ipsilesional SLF.^90^ The current study is the first to show that myelin within the SLF is disrupted after stroke but may have the capacity to recover after a large dose of skilled motor practice. Future studies should probe the SLF in addition to motor regions, to advance our understanding of recovery from stroke.

In contrast to the pattern of change noted in the SLF, MWF in the PLIC was not altered by skilled motor practice in the current study. Given the large dose of movement practice delivered in our study, this was a surprising result. However, it may be that the lack of change noted in the current study shows that MWF in the PLIC for both groups was sufficient to meet the demands of our experimental task. Future work could include larger sample of individuals with varying degrees of function to explore the role of PLIC MWF following skilled training in chronic stroke.

### Limitations

Several limitations may influence interpretations of the observed findings. First, we considered individuals with a wide range of motor impairment and functional capacity after stroke. Despite this variability, all participants showed motor learning related change in behaviour as indexed by decreasing (faster) movement times over the course of training. In addition, inclusion of individuals with varied levels of motor impairment also allows our findings to apply to a wide range of individuals with stroke. Future studies should evaluate sub-groups of individuals with similar motor impairment, functional ability or lesion location to increase understanding of implications to stroke recovery. Second, inclusion of a control group of individuals who did not engage in motor practice but were scanned four weeks apart would help rule out the possibility that changes in MWF were a result of reliability issues. However, MWF has shown excellent test re-test reliability and suitability for longitudinal studies.^91^ Therefore, it seems unlikely that the changes observed were a product of reliability issues. Third, our conclusions cannot be extended to individuals with acute stroke as we only studied individuals in the chronic stage of recovery. This experimental approach instills confidence that the observed changed were a result of motor training and not associated with spontaneous recovery. It will be important to consider patterns of myelin plasticity related to different interventions in the acute and sub-acute stages of recovery from stroke. Fourth, we were limited by the time frame of our MRI scanning. At present we do not understand precisely when change in MWF occurred. It is also possible that individuals change MWF at different points in practice which could help explain why we did not observe change in MWF in controls. It will be important for future work to assess the time course of change in myelin across practice in all individuals. These efforts may allow for the timing and the amount of practice to be personalized to optimize myelin plasticity and behavioural change in individuals with stroke.

### Conclusion

The current study shows that myelin in the human brain is neuroplastic and supports motor learning after stroke. Myelin water fraction was increased by a high dose of skilled motor practice in individuals with chronic stroke as compared to healthy controls, and associated with motor learning. We provide evidence that baseline myelination of motor and visuomotor regions correlate with motor learning rate in individuals with stroke. Ultimately, our findings suggest that individuals with stroke relied on change in a broad array of white matter regions that support functions that are important for learning and include areas that support visuomotor integration, cognition, and attention. In sum, myelin plasticity appears to be important for motor learning in the stroke-injured brain.

## Abbreviations

CP: Cerebral peduncles
CCG: Cingulum-cingulate gyrus
COV: Coefficient of variation
CST: Corticospinal tract
DTI: Diffusion tensor imaging
FA: Fractional anisotropy
FM-UE: Fugl-Meyer upper extremity scale
GRASE: Gradient- and spin-echo
MNI: Montreal Neurological Institute coordinate system
MCRI: Multi-component relaxation imaging
MWF: Myelin water fraction
PCR: Posterior corona radiata
PLIC: Posterior limb of internal capsule
PTR: Posterior thalamic radiations
ROI: Region of interest
SCR: Superior corona radiata
SLF: Superior longitudinal fasciculus

## Data availability

Source data for main figures (Figures 3-5 and Supplementary Figure 1, excel files) are included.

---

**Supplementary Figure 1.**
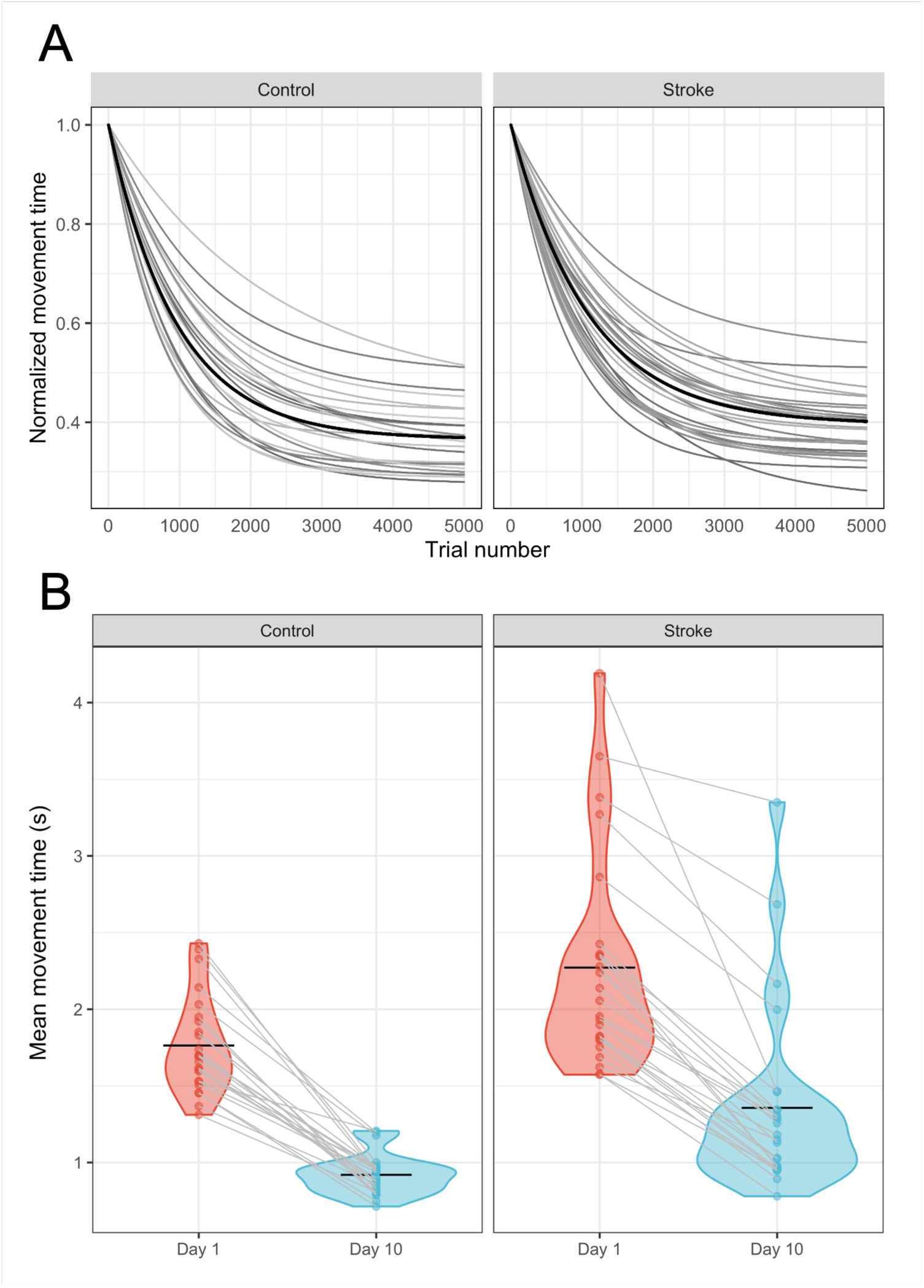
Movement times. (**A**) Normalized movement times for each individual participant and group mean (black line) across all trials. Decreased (faster) movement times over the course of training occurred in both groups. (**B**) Raw movement time of Day 1 (trial 1-500) and Day 10 (trial 4501-5000) and participant means (black line).

**Supplementary Table 1.** Correlation coefficients for the relationship between learning rate (alpha) and pre-training MWF per ROI and group.

| ROI | Stroke |  | Control |  |
| --- | --- | --- | --- | --- |
|  | *r*-value | *P*-value | *r*-value | *P*-value |
| Cerebral peduncles | 0.50 | 0.01* | -0.12 | 0.58 |
| Superior corona radiata | 0.50 | 0.02* | 0.38 | 0.07 |
| Posterior limb internal capsule | 0.43 | 0.03* | 0.49 | 0.02* |
| Posterior thalamic radiations | 0.18 | 0.38 | 0.12 | 0.58 |
| Posterior corona radiata | 0.50 | 0.01* | 0.24 | 0.27 |
| Cingulum-cingulate gyrus | 0.05 | 0.83 | 0.37 | 0.07 |
| Superior longitudinal fasciculus | 0.19 | 0.36 | 0.2 | 0.35 |

\* $P < 0.05$

